# Progressive impairment of directional and spatially precise trajectories by TgF344-AD Rats in the Morris Water Task

**DOI:** 10.1101/282392

**Authors:** Laura E. Berkowitz, Ryan E. Harvey, Emma Drake, Shannon M. Thompson, Benjamin J. Clark

**Author notes:** To whom correspondence should be addressed: Benjamin J Clark, Ph.D.

## Abstract

Spatial navigation is impaired in early stages of Alzheimer’s disease (AD), and may be a defining behavioral marker of preclinical AD. Nevertheless, limitations of diagnostic criteria for AD and within animal models of AD make characterization of preclinical AD difficult. A new rat model (TgF344-AD) of AD overcomes many of these limitations, though spatial navigation has not been comprehensively assessed. Using the hidden and cued platform variants of the Morris water task, a longitudinal assessment of spatial navigation was conducted on TgF344-AD (n=16) and Fischer 344 (n=12) male and female rats at three age ranges: 4 to 5 months, 7 to 8, and 10 to 11 months of age. TgF344-AD rats exhibited largely intact navigation at 4-5 and 7-8 months of age, with deficits in the hidden platform task emerging at 10-11 months of age. In general, TgF344-AD rats displayed less accurate swim trajectories to the platform and a wider search area around the platform region compared to wildtype rats. Impaired navigation occurred in the absence of deficits in acquiring the procedural task demands or navigation to the cued platform location. Together, the results indicate that TgF344-AD rats exhibit comparable deficits to those found in individuals in the early stages of AD.

## Introduction

Alzheimer’s disease (AD) is the most common form of dementia in the United States and is characterized by progressive cognitive decline and neurodegeneration (Association and others, 2017). AD pathology, marked by amyloid plaques and neurofibrillary tangles that accumulate throughout the limbic system and hippocampus, is predicted to initiate up to 20 years prior to the onset of behavioral symptoms (Gao et al., 1998; Mielke et al., 2014). Although the long preclinical period poses a significant challenge to early diagnosis of AD (Dubois et al., 2016; Graham et al., 2017) subtle changes in memory, as observed in amnestic mild cognitive impairment (aMCI), can indicate an increased risk of progressing to dementia (Petersen, 2004; Winblad et al., 2004). However, because not all cases of aMCI progress to AD, there is a growing need for sensitive behavioral assessments for AD diagnosis.

Mounting evidence suggests that spatial disorientation, sometimes referred to as wandering, are among the earliest memory complaints in AD (Bianchini et al., 2014; Bird et al., 2010; Chan et al., 2016; Guariglia and Nitrini, 2009; Pai and Jacobs, 2004; Yew et al., 2013). In general, disorientation is characterized as deficient localization of hidden goals (Hort et al., 2007), a loss of direction sense (Cushman et al., 2008; deIpolyi et al., 2007; Monacelli et al., 2003; Tu et al., 2015), or an impairment in correctly identifying familiar spatial scenes after a small change in view-point (Bird et al., 2010; Chan et al., 2016). Deficits in establishing or utilizing map-like (allocentric) frameworks for navigation are frequently linked with the earliest stages of AD, while later stages are associated with deficits in simpler forms of navigation, such as approaching cues, or utilizing egocentric movements to guide behavior (Hort et al., 2007). Navigation can therefore serve as an early marker of AD, with some studies indicating that disoriented patients are more likely to convert from aMCI to an AD diagnosis (Bird et al., 2010; Laczó et al., 2009).

Animal models have been utilized to better understand spatial disorientation in AD, however most fail to produce the full spectrum of amyloid and tau dysfunction (Do Carmo and Cuello, 2013; LaFerla and Green, 2012). A recent rat model of AD, called the TgF344-AD rat, has been developed to express both the Swedish form of mutant amyloid precursor protein gene and human mutant presenilin 1 and progressively develop a comprehensive profile of AD pathology (Cohen et al., 2013). Importantly, TgF344-AD rats display characteristic plaque and tangle pathogenesis, as well as neuroinflammation, and neurovascular dysfunction resulting in substantial cell loss (Joo et al., 2017). Elevated amyloid, tau, and activated microglia pathology can be detected within both the entorhinal cortex and hippocampus at 6 months of age, and by 16 months of age, the expression of amyloid and tau increase substantially (Cohen et al., 2013; Rorabaugh et al., 2017). A recent study has reported that reductions in the strength of excitatory transmission can be detected at entorhinal-to-dentate gyrus synapses as early as 6 months (Smith and McMahon, 2018). These synaptic changes precede reductions in neurotransmission at CA3-CA1 hippocampal synapses, which begin around 9 months of age. This emerging pattern of pathogenesis and altered signaling along the Trisynaptic circuit is consistent with the human condition (Braak and Braak, 1995; Thal et al., 2002).

Currently, only a small number of studies have investigated the time-course of spatial navigation impairment in the TgF344-AD rat model (Cohen et al., 2013; Pentkowski et al., 2017; Rorabaugh et al., 2017). While experiments have determined that spatial deficits can be detected after 15 months of age (Cohen et al., 2013; Rorabaugh et al., 2017; Voorhees et al., 2017), the onset of navigation impairment has been less clear. Some studies have reported that TgF344-AD rats are largely unimpaired in navigating to a fixed hidden goal location between 4 and 6 months of age (Cohen et al., 2013; Pentkowski et al., 2017), while others have reported deficits at 6 months of age (Rorabaugh et al., 2017). The variability between studies could be related to subtle differences in performance by male and female TgF344-AD rats, which are often pooled into a single group. Sex differences in spatial behavior in AD have been reported in other rodent models of AD (Clinton et al., 2007; Granger et al., 2016; Mielke et al., 2014), and is additionally supported by the fact that females exhibit a disproportionally higher rate of progression to AD and increased severity of clinical dementia. Further, given the cross-sectional design of previous studies, and the fact that TgF344-AD display differing locomotor behaviors as early as 6 months of age (Cohen et al., 2013), non-spatial variables associated with procedural learning could also play a role.

The overall aim of the present study was to characterize spatial navigation in TgF344-AD rats at early stages of development. A longitudinal design was employed to capture compensatory behaviors possibly secondary to burgeoning pathology (Morganti et al., 2013) and to control for age-effects commonly attributed with procedural task demands (van Groen et al., 2002; Vicens et al., 2002). Both male and female subjects were tested at three age ranges starting from 4 to 5 months, 7 to 8 months, and 10 to 11 months in the hidden and cued platform variants of the Morris water task (Harker and Whishaw, 2002; Morris, 1984; Pentkowski et al., 2017; Vorhees and Williams, 2006). Several standard measures were used to describe the performance and spatial distribution of movements in the pool, including escape latency, path length, and platform proximity. However we also provided an assessment of directional disorientation by quantifying the proportion of direct swims toward the platform region at each age of testing (Garthe et al., 2009; Gehring et al., 2015). Finally, a convolution analysis was employed to determine how switching to direct swims or becoming more efficient at less direct swims contributes to task performance. Consistent with what has been reported in human AD, we hypothesized that rats would express age-dependent impairments in the accuracy of their swim trajectories in the hidden platform variant of the Morris water task. We additionally hypothesized that these impairments would be expressed independent of motor and procedural deficits, and co-occur with intact acquisition of cued platform learning.

## Methods

### Subjects

Sixteen TgF344-AD (Tg) rats counterbalanced for sex and expressing mutant human amyloid precursor protein (APPsw) and presenilin 1 (PS1ΔE9) were obtained directly from the Rat Resource & Research Center (Columbia, MO). Twelve wild type Fischer 344 (WT) rats (Harlan laboratories, Indianapolis, IN) counterbalanced for sex served as control subjects. Subjects were maintained under controlled temperature (21 ± 2 ◦C), were housed with Tg or WT pairs, and were kept on a 12 hour light/dark cycle (lights off at 09:00 AM). Food and water was provided *ad libitum* throughout the duration of the study. Animal care practices and experiments were approved by the University of New Mexico Institutional Animal Care and Use Committee and adhered to the APA ethical principles of animal use.

### Longitudinal Design

A longitudinal mixed design was used to assess changes in navigation in the hidden and cued platform variants of the Morris water tasks at three age time-points. Subjects were placed into groups by genotype (Tg or WT) and further grouped by sex (male or female). At the start of testing, subjects’ age ranged from 4.30-5.7, 7.5-8.9, and 10.3-11.7 months at each respective time point. For clarity, we refer to these ages throughout the remaining manuscript as, 4-5 months, 7-8 months, and 10-11 months of age. Tasks were administered in the same temporal order at each time point (Fig. 1A), whereby the hidden platform task was administered before cued platform training. Procedures were maintained between time points and experimenters did not change throughout the duration of the study. All rats were naïve to experimentation prior to the start of the first experiments at 4 months of age.

**Figure 1.**
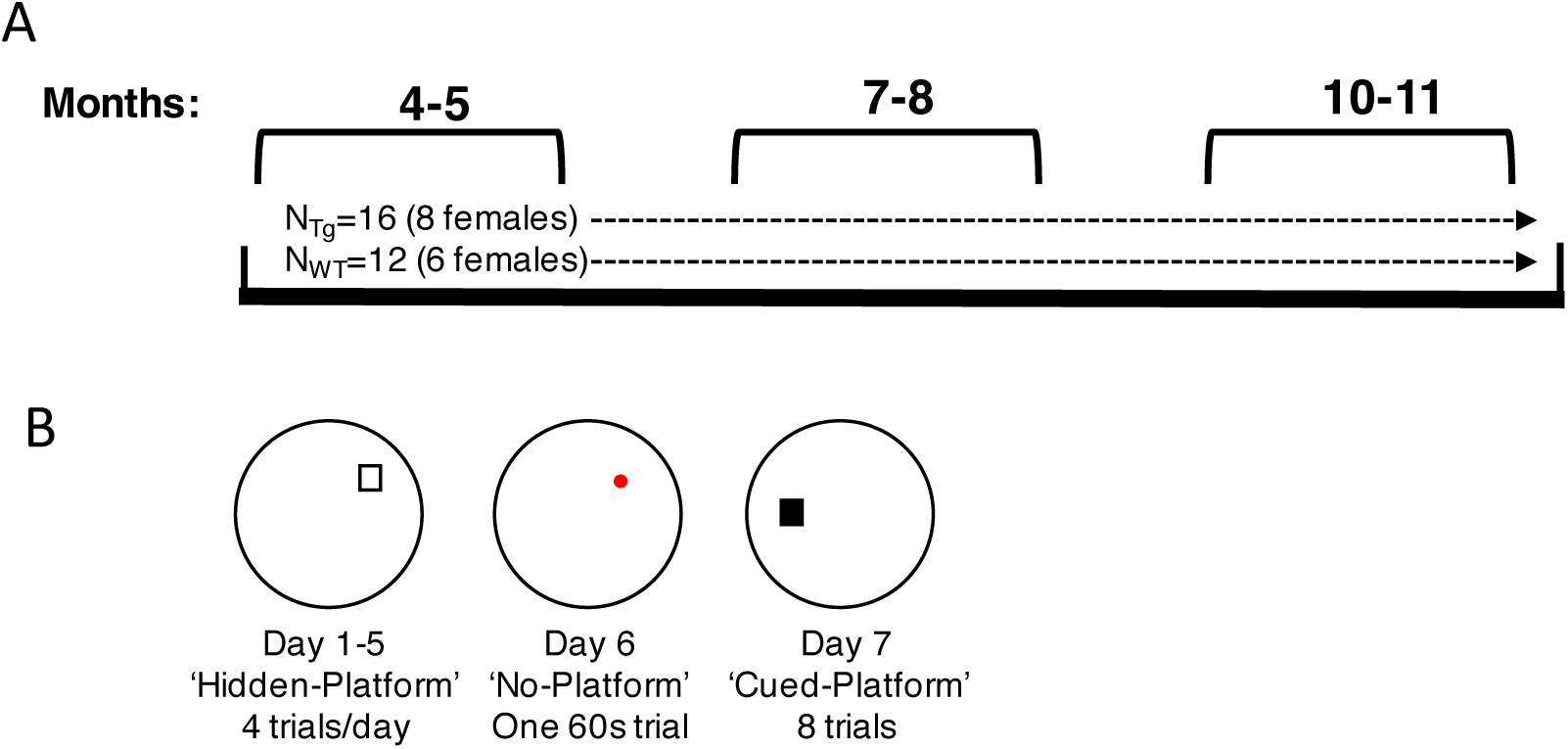
Study Time line. (A) TgF344-AD (n=16) and wild-type F344 rats (n=12) underwent testing at three age points (shown in months). (B) The temporal order of testing consisted of 5 days of training followed by a no-platform probe trial and finally a cued-platform test on day 7. White square=hidden platform, red circle=prior platform area, black square= cued-platform.

### Apparatus and Testing Room

A circular pool (150 diameter, 48 cm high) with a white inner wall situated on a wooden frame (50 cm high) and a 16 cm x 16 cm plastic escape platform covered with a metal grate with a height of 25cm was used for all tasks. The pool was filled with water (20 - 22°C) until the level reached ∼2.5 cm above the top of the platform. Non-toxic white paint (∼500 ml) was used to make the water opaque. Distal environmental cues were composed of various objects (movie posters, thin particle board hangings) and furniture (desks & bookshelves) and were maintained in fixed locations during training at each time-point. However, the cues, cue layout, pool, and platform location was modified at each time-point (Fig. 1B). An overhead camera was fastened above the pool to record swim behavior for subsequent analysis.

### Morris Water Task - Hidden Platform

The purpose of the hidden platform variant of the Morris water task is to assess animal navigation to a precise spatial location based on multiple allocentric spatial cues particularly those associated with the features of the distal environment (Clark et al., 2015; Harker and Whishaw, 2002; Morris, 1984; Vorhees and Williams, 2014). Rats were given 4 trials per day for 5 consecutive days. An individual trial was composed of randomly placing the rat into the pool at one of four equidistant drop locations facing the wall. Once the platform was found, the subject remained on the platform for 10 seconds. Drop location varied between time point/trials/days, but was maintained between subjects. At the end of the trial, the rat was returned to a holding cage, during which time the remaining rats in the cohort were tested (10-20min duration). Subjects were run in groups of seven for each task. At the end of the 4 trials, rats were returned to their home cages in the colony room and the same procedure was repeated the next day. Time to reach the platform was recorded by an experimenter at the end of each trial. The mean escape latencies (seconds) for each animal were calculated for each day (every 4 trials).

### No-Platform Probe Test

After 5 days of training in the hidden platform task, a probe trial was conducted in which the platform was removed from the swimming pool. The probe trial was conducted 24 hours after the final trial of the fifth training day. The probe trial was composed of releasing rats from the side of the pool opposite the platform. Rats were permitted to swim for a total of 60 seconds. After the probe test, rats were removed from the pool and placed in a holding cage.

### Morris Water Task - Cued Platform

A cued-platform test was performed to verify that animals could navigate toward a cue directly associated with the platform location (Clark et al., 2013; Clark and Taube, 2009; Morris, 1984). In this task, a 10 cm diameter black ball with a white horizontal stripe was attached to a metal rebar and placed above the center of the platform (11.5 cm above the water). The cue was visible from any location in the pool. Subjects were trained for a single day consisting of 8 trials. Again, a trial consisted of placing a rat by hand into the water facing the wall of the pool at 1 of the 4 starting positions (trial length 60 sec; time on platform 10 sec). If the platform was not found within 60 seconds, the experimenter guided the subject to the platform by hand. Time to reach the platform was recorded by an experimenter at the end of each trial.

### Behavioral Analysis

From the video records, behavioral coders blind to experimental group manually tracked each animal’s location in the pool. Tracking and analysis was performed for each trial of the hidden platform, no-platform probe, and cued platform experiments (Clark et al., 2015; Pentkowski et al., 2018). First, raw video files were converted to JPEG images in a Linux bash shell using FFMPEG (https://www.ffmpeg.org). Image files were then imported into Fiji (https://imagej.nih.gov/ij/) and x- y- coordinates of the animal’s nose was acquired for each video frame (10 frames/sec) using the Manual Tracking plugin. Custom Matlab (R2017, The MathWorks, Natick, MA) scripts were designed to smooth the tracked swim paths using the runline function from the Chronux toolbox (www.chronux.org). The smoothed paths were then analyzed for path length (cm), swim speed (cm/sec), platform proximity (cm), and search area. Platform proximity was measured by summing the distance between the subject’s location and the center of the platform location over 1 sec intervals (Gallagher et al., 1993). The distance from the drop location to the center of the platform was then subtracted from the total summed distances as drop locations were variably distant to the platform. Search area was obtained by first dividing the pool surface into a matrix of 50 x 50 bins (each bin = 3cm x 3cm). Search area was expressed as the proportion of the pool visited by the animal (number of visited bins divided by the total number of bins in the matrix).

For no-platform probe trials, the pool was divided into four equal quadrants and the total dwell time in each quadrant was determined. From these measures, a platform preference score was calculated by taking the average difference in dwell time between the platform quadrant and the opposite quadrant (Harker and Whishaw, 2002). We also calculated an average platform proximity for the probe test by taking an average of the distance between the subject’s location and the center of the platform location across the 60 second probe test.

Additionally, we performed a detailed swim trajectory analysis to determine the number of directed swim movements toward the hidden platform during training (Brody and Holtzman, 2006; Garthe et al., 2009; Gehring et al., 2015; Graziano et al., 2003; Ruediger et al., 2012; Stone et al., 2011; Wolfer and Lipp, 2000) Briefly, a custom Matlab script automatically segmented whole paths into 200 cm increments that overlapped 70% to minimize labelling bias secondary to segment start/end points in order to capture the use of various movement types within a given trial (Gehring et al., 2015). An expert coder blinded to group manually categorized the path segments into 11 movement subtypes based on previous work (Garthe et al., 2009; Gehring et al., 2015). For clarity of analysis, the movements were further divided into 4 broad categories. Figure 4 provides a description of each movement subtype and analysis category. Briefly, target-direct movements included trajectories that contained no looping and path lengths less than 200cm and were further defined as a direct movement to platform (i.e. direct) or meandering/arched movements towards platform (i.e. circuitous-direct). Target-indirect movements included circling or searching next to the platform (i.e. target-search), and trajectories that passed by platform (i.e. target-scanning). Spatial-indirect movements involved patterns indicative of spatial processing but were not directed towards the platform location, such as maintaining a swim path set approximately the distance of the platform to the wall (i.e. chaining), searching of a focal area in the pool (i.e. focused-search), sustained swimming in the center of the pool (scanning), and sweeping swim paths traversing pool quadrants (i.e. scanning-surrounding). Non-Spatial movements including wall hugging (i.e. thigmotaxis), using the wall to head in a random trajectory away from the pool wall (i.e. incursion), and a path that circle over itself (i.e. looping). If a segment displayed characteristics of multiple movement types, the path was labeled with all applicable types.

Lastly, an experimenter blind to group quantified the number of non-spatial learning errors during training trials for the hidden platform experiment. As previously described (Harker and Whishaw, 2002; Saucier et al., 1996; Saucier and Cain, 1995), non-spatial errors included diving behavior (diving below the surface of the water during a trial), floating (the absence of swimming for more than 3 sec), platform deflections (the failure to obtain purchase onto the platform after contact), mounting errors (the failure to climb the platform after 1 sec of contacting the platform), and jumping off the platform. Each error was assigned a score of 1 and a total was obtained by summing the errors (Harker and Whishaw, 2002)

### Statistical Analysis

Statistical analyses were performed using SPSS version 26 (IBM, Armonk, NY) and MATLAB. Mixed model ANOVAs were used to assess differences within or between time points. Between-subject factors consisted of sex (female, male) and genotype (Tg, WT). Outcome measures were evaluated in blocks of four trials (day 1, day 2, day 3, day 4, day 5) or time-point. Log transformations were applied to data that did not meet the assumption of normality. A Greenhouse-Geisser correction was applied to data that did not meet the assumption of sphericity. Two-way ANOVAs were used to compare performance on the cued platform task within a time point using genotype and sex as between subject factors. The results of the detailed path analyses were subjected to Chi square tests. Robust linear regressions using a Tukey’s bisquare weight function to minimize the effect of outliers were used to determine the proportion of variance of task performance accounted for by convolved factors (described further in the Results section). Both significant (p <.05) and trending non-significant (n.s.) (.1>p>.05) results are reported.

## Results

One subject was excluded from the analyses at 10-11 months of age after developing glaucoma. Furthermore, one subject died during testing at 10-11 months of age and thus was not included in the analysis below.

### Hidden Platform Task

#### Swim Latency

Figure 2A displays measures of escape latency for groups at each time-point. On average, animals in all groups showed a progressive decrease in escape latency at each age of testing, indicating that subjects could learn the location of the platform. This observation is supported by a significant day effect for all groups at the three time-points (*Fs*≥ 15.25, *ps*≤.001). While there were no significant genotype or sex differences at 4-5 and 7-8 months of age, swim duration by Tg rats at 10-11 months of age was appreciably greater than WT animals. This was confirmed by a significant genotype difference in escape latency at the final time-point (*F*(1, 23)=5.76, *p*=.025). In addition, at 10-11 months of age, there was a trend in which Tg females had greater escape latencies relative to all other groups (≤8.1s, ± ≤1.1 sec; *F*(1, 23)=3.10, *p*=.092).

**Figure 2.**
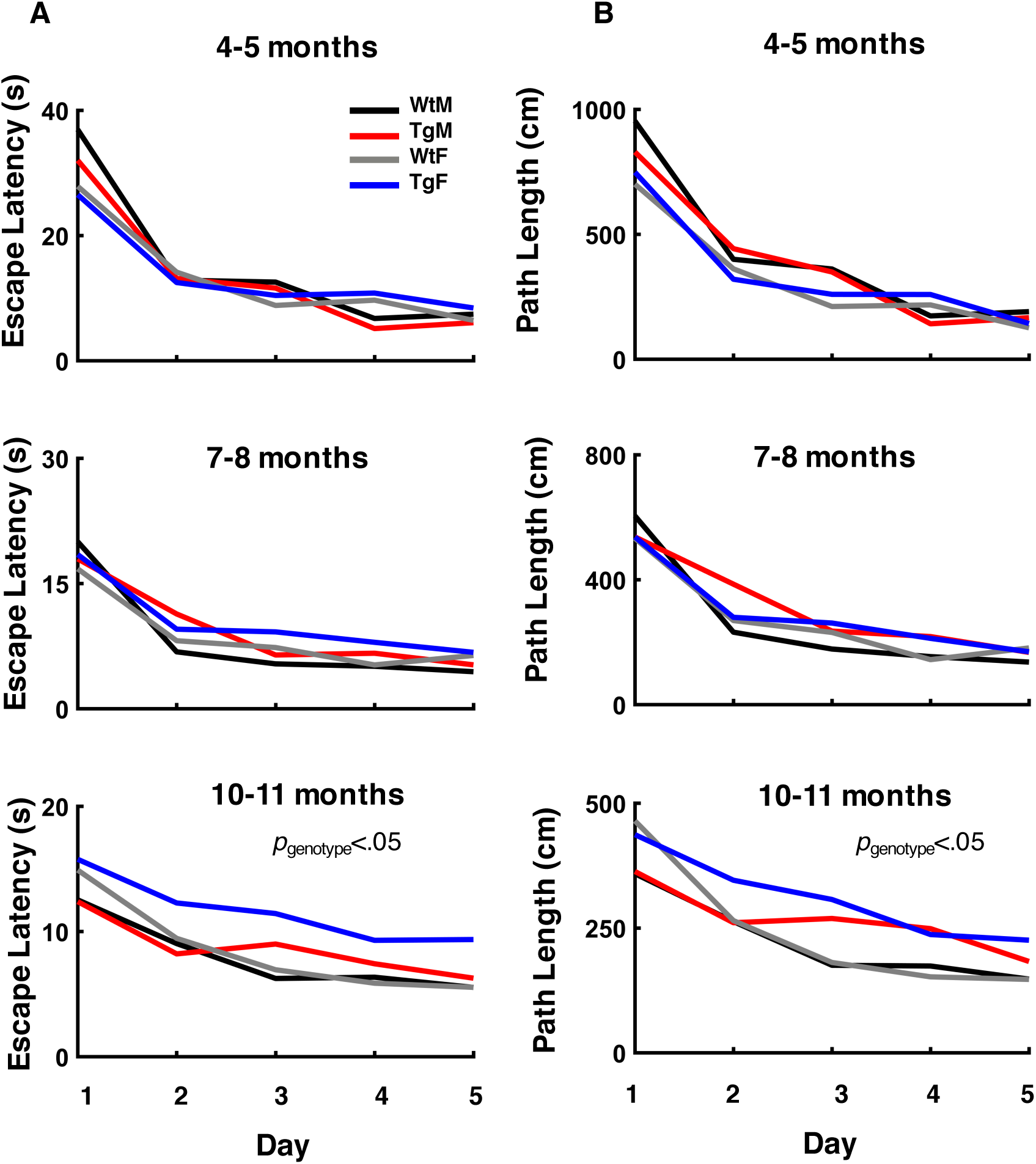
TgF344-AD rats took more time to reach the platform and had longer path lengths during hidden-platform training compared to WT controls starting at 10-11 months of age. (A) Mean escape Latency in seconds plotted over days at all age points. Main effect of genotype observed at 10-11months, (*p*<.05, Mixed ANOVA). (B) Mean path length in centimeters plotted over days at all age points. Main effect of genotype observed at 10-11months, (*p*<.05, Mixed ANOVA). Groups distinguished by color: black=Wild Type males (WtM), red=Transgenic males (TgM), gray=Wild Type females (WtF), blue=Transgenic females(TgF).

#### Path Length

Measures of path length mirrored that of escape latency in showing that animals reduced the length of their swims during training at each time-point (Fig. 2B). Again, a mixed ANOVA indicated significant day effects for all groups at the three time-points (*Fs*≥ 15.01, *ps*≤.001). Sex differences in swim length were detected at 4-5 months of age (*F*1, 24) =4.69, *p*=.040), but not at 7-8 or 10-11 months of age. Although there were no genotype differences at 4-5 and 7-8 months of age, Tg rats had significantly longer paths by 10-11 months (*F* (1, 23)=5.18, *p*=.032).

#### Spatial Distribution of Swim Paths

The greater path length displayed by Tg rats at 10-11 months of age may reflect a pattern of behavior in which movements are restricted to specific non-platform locations or distributed over wide areas of the pool. To better understand the spatial distribution of swim paths during hidden platform training, we analyzed the overall search area and the relative proximity to the platform of swim paths during each training trial (Fig. 3). ANOVAs conducted on search area and platform proximity indicated a significant day effect at each age (*Fs*≥ 30.72, *ps*≤.001), however there was no evidence of a significant genotype or sex difference at 4-5 or 7-8 months of age. Nonetheless, Tg rats displayed wider search distances relative to the platform at 10-11 months (Fig. 3A). The observation is supported by significant genotype (*F* (1, 23)=5.75, *p*=.025) and genotype by day effects (*F*(4, 92)=2.58, *p*=.043). Similarly, Tg rats displayed a search area that was wider compared to WT animals at 10-11 months of age (Fig. 3B). This difference was confirmed by a significant ANOVA for genotype at 10-11 months (*F*(1, 23)=4.69, *p*=.041). Collectively, these findings indicate that the swim behavior of Tg rats was less confined to the platform region and distributed to wider regions of the pool.

**Figure 3.**
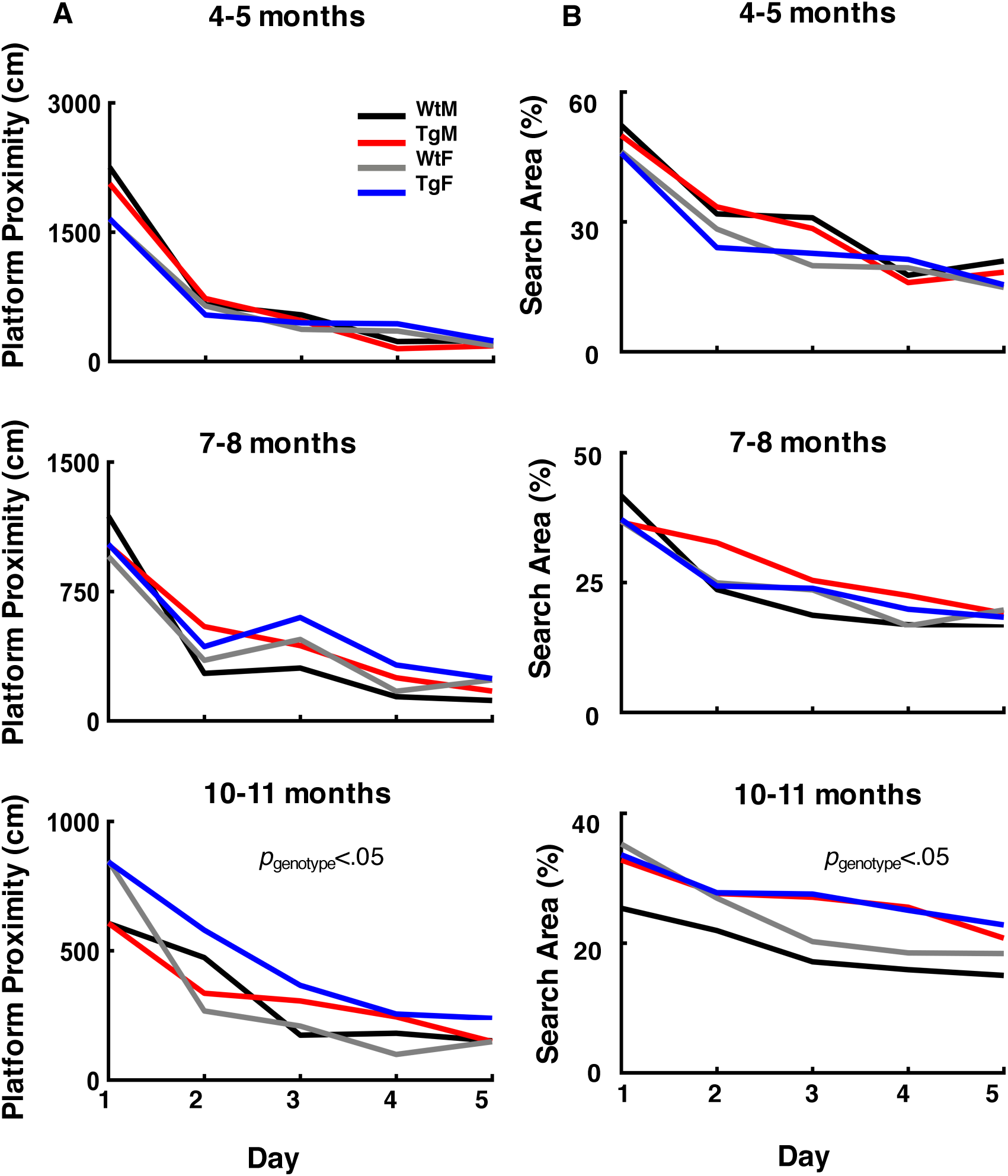
During hidden-platform training, TgF344-AD rats swam further from the platform and swam a larger proportion of the pool area relative to WT controls starting at 10-11 months of age. (A) Mean platform proximity in centimeters plotted over days at all age points. Main effect of genotype observed at 10-11months, (*p*<.05, Mixed ANOVA). (B) Percentage of search area explored plotted over days at all age points. Main effect of genotype observed at 10-11months, (*p*<.05, Mixed ANOVA). Groups distinguished by color: black=Wild Type males (WtM), red=Transgenic males (TgM), gray=Wild Type females (WtF), blue=Transgenic females (TgF).

#### Swim Trajectory Analysis

In well trained control animals, swim trajectories are typically composed of movements characterized by direct paths beginning at the start point and ending at the platform location. The findings summarized in the sections above suggest that 10-11-month-old Tg rats may express fewer of these directed swims. To address this possibility, we classified swim paths into 11 movement categories (Fig. 4). These swim movements were then merged into 4 categories based on their directedness to the platform (target-direct), a lack of directedness but proximity to the platform (target-indirect), or a tendency to organize movements in non-platform locations (spatial-indirect), or in a random search (non-spatial).

**Figure 4.**
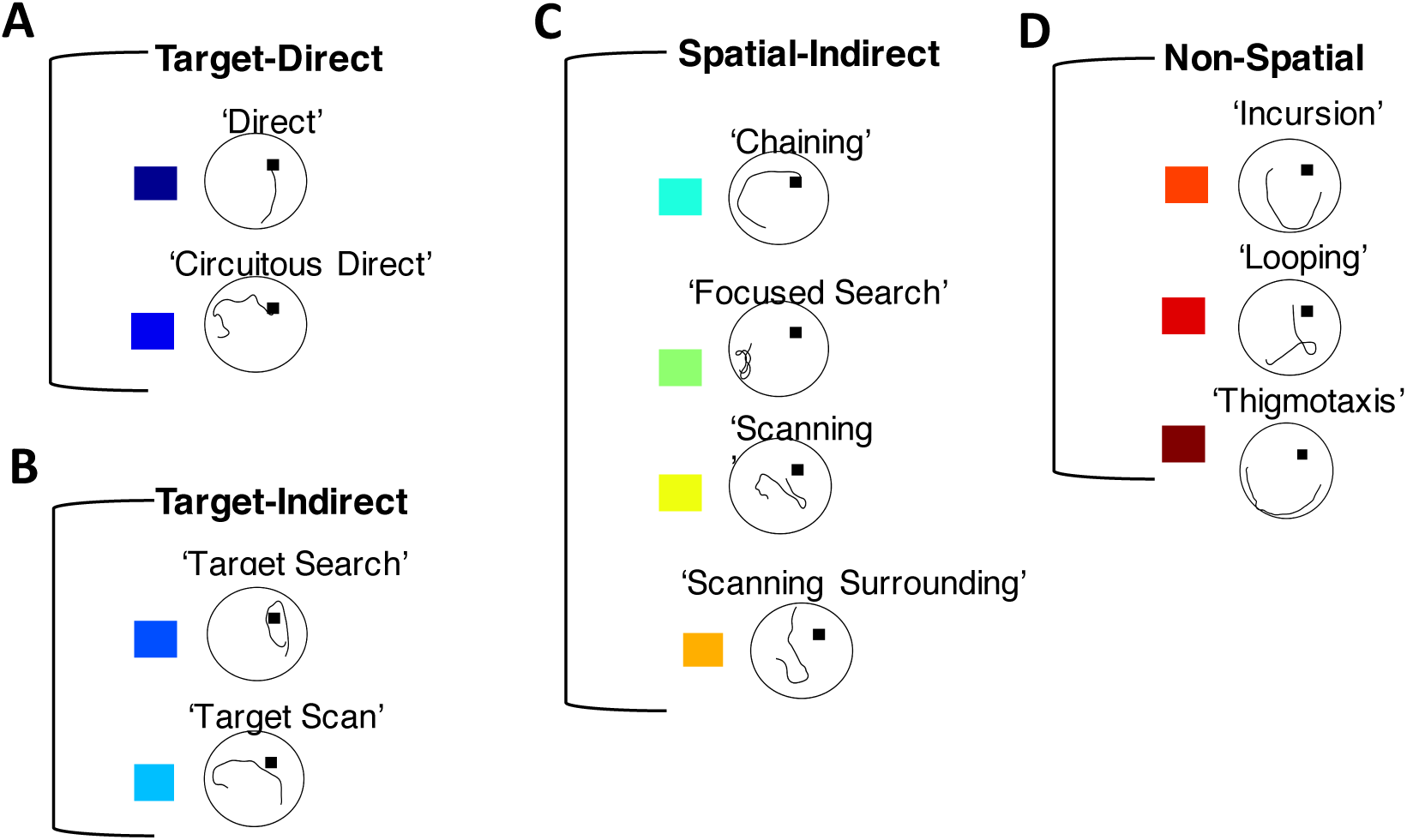
Representative examples of individual swim segments. (A) Target-direct (platform found within first segment). (B) Target-indirect (area around platform searched). (C) Spatial-indirect (spatial components not directed at platform). (D) Non-spatial (limited spatial components). See *Methods* for operational definitions of individual strategies.

Figure 5 and 6 show the proportion of each movement category for groups across the three experimental ages and testing days. As expected, both Tg and WT rats displayed an increasing number of direct trajectories as a function of training day at each age (Fig. 6A). However, the overall proportion of target-direct trajectories by Tg rats was significantly lower at 10-11 months of age (*X^2^(1) =* 12.43, *p*<.001). Interestingly, the proportion of direct trajectories by female Tg rats was appreciably lower compared to all other groups (*X^2^s ≥*5.96, ps≤.014). Additionally, inspection of target-direct trajectories at earlier ages suggest that Tg rats made fewer of these movements relative to WT animals. Indeed, group comparisons reached significance at 7-8 months of age (*X^2^(1) =* 5.82, *p*=.015), and although group differences were not significantly different at 4-5 months of age, Tg females made fewer target-direct trajectories compared to WT females (*X^2^(1) =* 4.07, *p*=.043). Together, these results support the conclusion that Tg rats make less directionally precise movements at 7-8 and 10-11 months of age, with a significantly lower proportion of these movements by female Tg rats.

**Figure 5.**
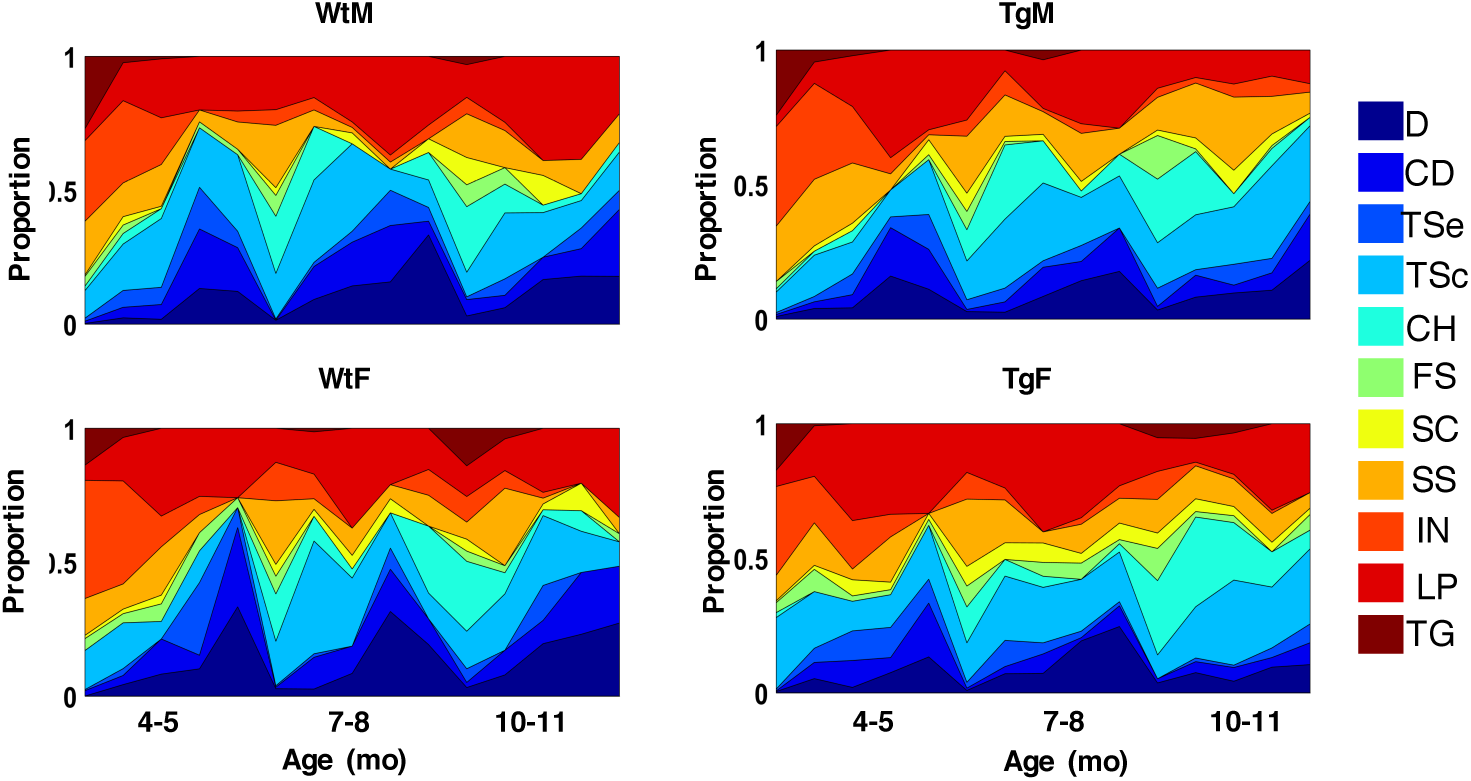
Relative proportion of swim strategies over days at each age of testing. Area plots showing the proportion of all strategies across days and age points. Colors indicate strategies, D=Direct, CD=Circuitous Direct, TSe=Target Search, TSc=Target Scan, CH=Chaining, FS=Focused Search, SC=Scanning, SS=Scanning Surroundings, IN=Incursion, LP=Looping, TG=Thigmotaxis. Note that Tg animals show fewer direct (D) and circuitous-direct (CD) compared to WT at 7-8 months and 10-11 months of age.

**Figure 6.**
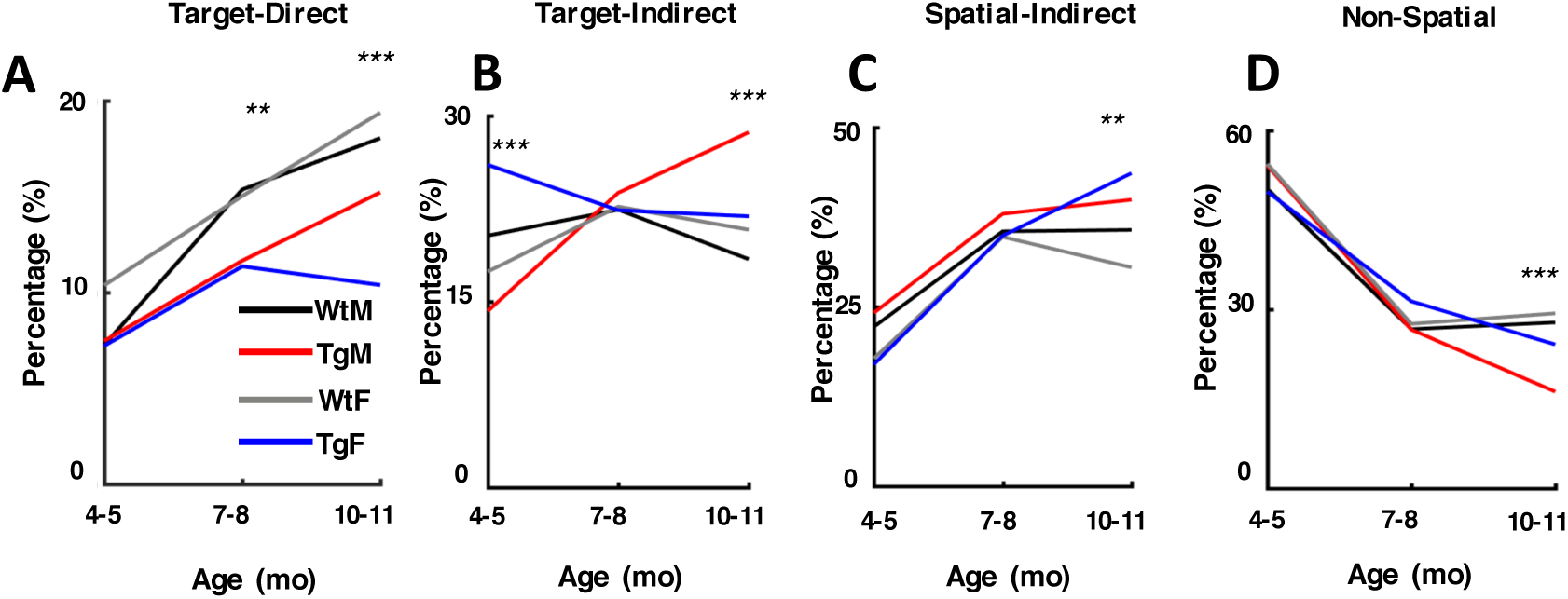
Line plots indicating the percentage of Target-Direct (A), Target-Indirect (B), Spatial-Indirect (C) and Non-Spatial (D) categories across ages. Note that Tg animals show a lower percentage of Target-Direct strategies starting at 7-8mo of age. Tg Males have a significantly higher proportion of Target-Indirect strategies at 10-11mo while Tg females use Spatial-Indirect strategies to a greater extent than WT females at 10-11mo. Tg animals show less Non-Spatial swim strategies compared to WT at 10-11 months (\**p*<.05; \*\**p*<.01; \*\*\**p*<.001).

The reduced frequency of target-direct paths by Tg rats suggest that other movements, including those directed near the platform (target-indirect) or in other non-platform locations (spatial-indirect and non-spatial), may be favored by Tg animals (Fig. 6B-D). Consistent with this hypothesis, Tg rats were found to express a greater proportion of target-indirect movements at 10-11 months of age (*X^2^(1) =* 5.93, *p*=.015). Furthermore, with respect to target-indirect movements, there was a clear sex difference with Tg males showing a significantly greater proportion of these paths relative to all other groups (*X^2^s*≥6.75, *ps*≤.009). Interestingly, this finding was apparent at 4-5 months, however; at that age Tg females had a greater proportion of target-indirect paths relative to all other groups (*X^2^s*≥6.82, *ps*≤.009). Additionally, Tg rats performed a greater number of spatial-indirect trajectories (*X^2^(1) =* 14.31, *p*<.001) and fewer non-spatial movements at 10-11 months (*X^2^*(1)=14.61, *p*≤.001). In sum, the reduced frequency of direct trajectories by Tg rats at 10-11 months of age corresponds with an increase in spatially restricted swim paths near the platform location (target-indirect) or in other pool locations (spatial-indirect).

#### Convolution Analysis

Collectively, the results described in the previous sections indicate that while general performance measures decreased across testing days, group differences were detected at 10-11 months of age. Tg rats expressed fewer direct trajectories toward the platform location at 10-11 months, and at earlier testing ages, suggesting that they utilized other movements to navigate to the platform locations. In other words, it is possible that repeated spatial training and procedural learning may have allowed Tg animals to utilize less-direct strategies that result in similar performance on standard measures. For example, an animal may become better at estimating the distance of the platform to the pool wall (i.e. chaining) over time. Thus, this movement may result in progressively shorter paths to the platform.

Given Tg rats perform a lower proportion of directed movements (either target-direct or target-indirect movements) at 10-11 months, we tested the hypothesis that Tg rat performance is more highly explained by becoming more efficient at non-direct movements rather than switching between non-direct to direct movement categories. To evaluate this hypothesis we utilized a convolution analysis (Brody and Holtzman, 2006; Garthe et al., 2009), which determines whether a change in the frequency of movement subtypes, or a change in the efficiency of swim movements, predicts trial path length (Fig. 7). First, an estimate of the frequency of each movement was calculated by dividing the number of movements in each category by the total number of path segments (Fig. 7B). Second, we estimated the efficiency of a given movement by multiplying the frequency of that movement during a trial by the trial path length. Thus, efficiency scores for a given movement subtype reflects the contribution by that movement to trial path length. Two models allowed us to assess the power of changes in efficiency or changes in frequency of movement subtypes in predicting trial path length. Changes in efficiency (ΔEff) scores were derived by keeping frequency constant across trials whereas changes in frequency (ΔFreq) scores were derived by keeping efficiency constant across trials (Fig 7C). Overall values for efficiency (i.e. average efficiency across all 20 trials) and frequency (i.e. frequency of each strategy across all 20 trials) were used as constants. Predicted path length therefore reflects the contribution of the non-constant factor on the trial path length (Fig 7D).

**Figure 7.**
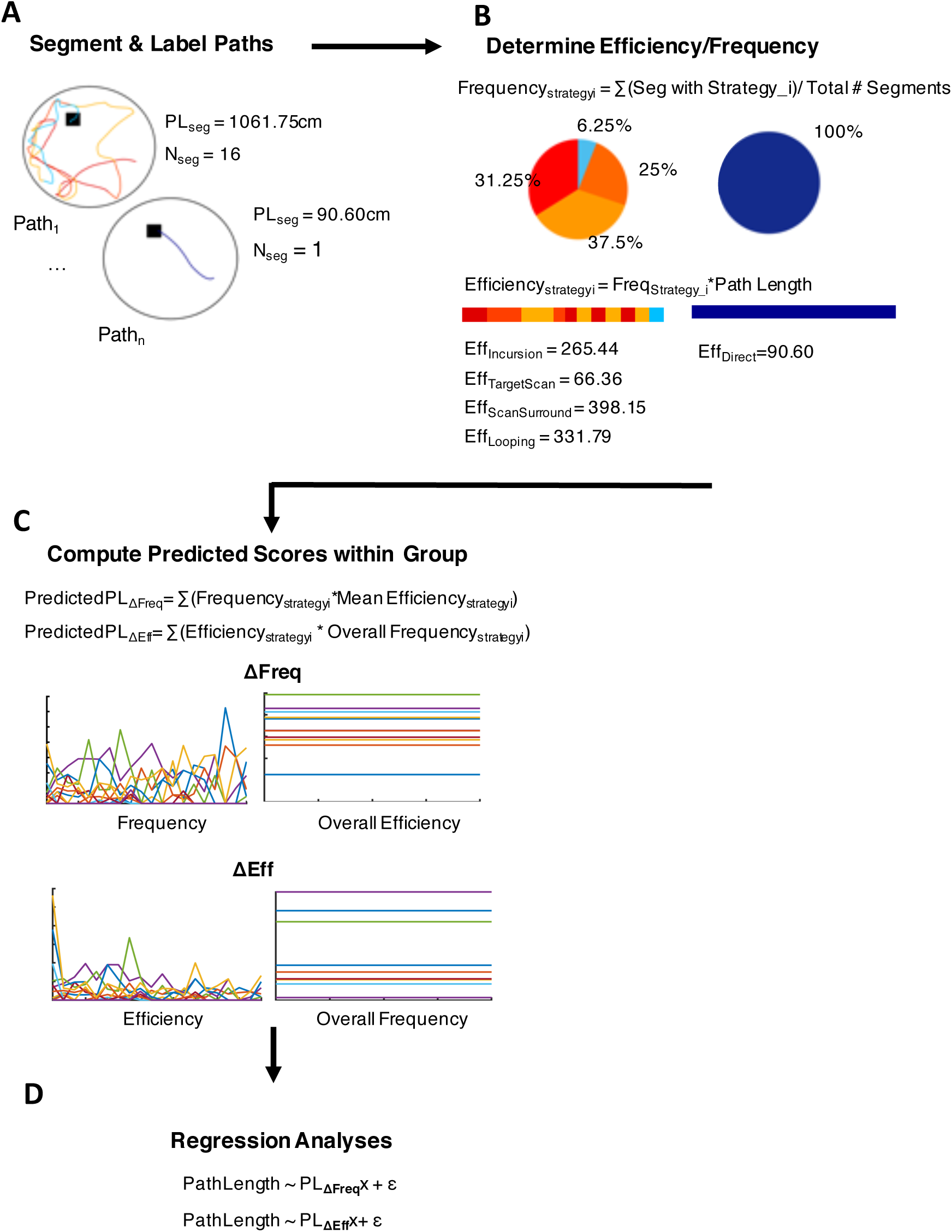
Analysis pipeline for assessment of swim movements using convolution analysis. (A) Example paths with movement category indicated by color, PL=path length, seg=segment. (B) Equations for computing the frequency and the efficiency of the segment and values for example paths denoted in (A). Pie charts represent the proportion of a given movement within a trial. Horizontal colored bar represents the proportion of a given movement segment over time across the relative distance. (C) Convolution analysis derived scores for frequency-fixed (Δeff represents changes in efficiency) and efficiency-fixed (Δfreq represents changes in frequency). (D) Model terms for linear regressions for each model.

Figure 8 summarizes the regression results which indicate that switching between movement categories and becoming efficient at swim movements is significantly predictive of trial path length for all animals. However, switching between movements takes up 27% less of the variance for Tg males and 16% less of the variance for Tg females relative to WT males and WT females, respectively (Fig. 9). Interestingly, Tg male and females exhibited differences in the predictive power of each factor. Specifically, at 10-11 months, switching between movements is a stronger predictor of performance for Tg males (β=4.12, t(18)=4.406*, p*<.001) relative to Tg females (β=1.77, t(18)=4.36*, p*<.001). Furthermore, although changes in efficiency take up a larger proportion of the variance for task performance in Tg females relative to Tg males (67% versus 45%, respectively), changes in efficiency hold less predictive weight for Tg females (β=.993, t(18)= 5.72*, p*<.001) relative to Tg males (β=1.80, t(18)= 3.67*, p*<.001). Overall, these results indicate that the performance of Tg animals is less associated with switching between movement categories than WT animals, supporting our hypothesis that Tg animals are more likely the improve the efficiency of their movements than switch to more direct trajectories.

**Figure 8.**
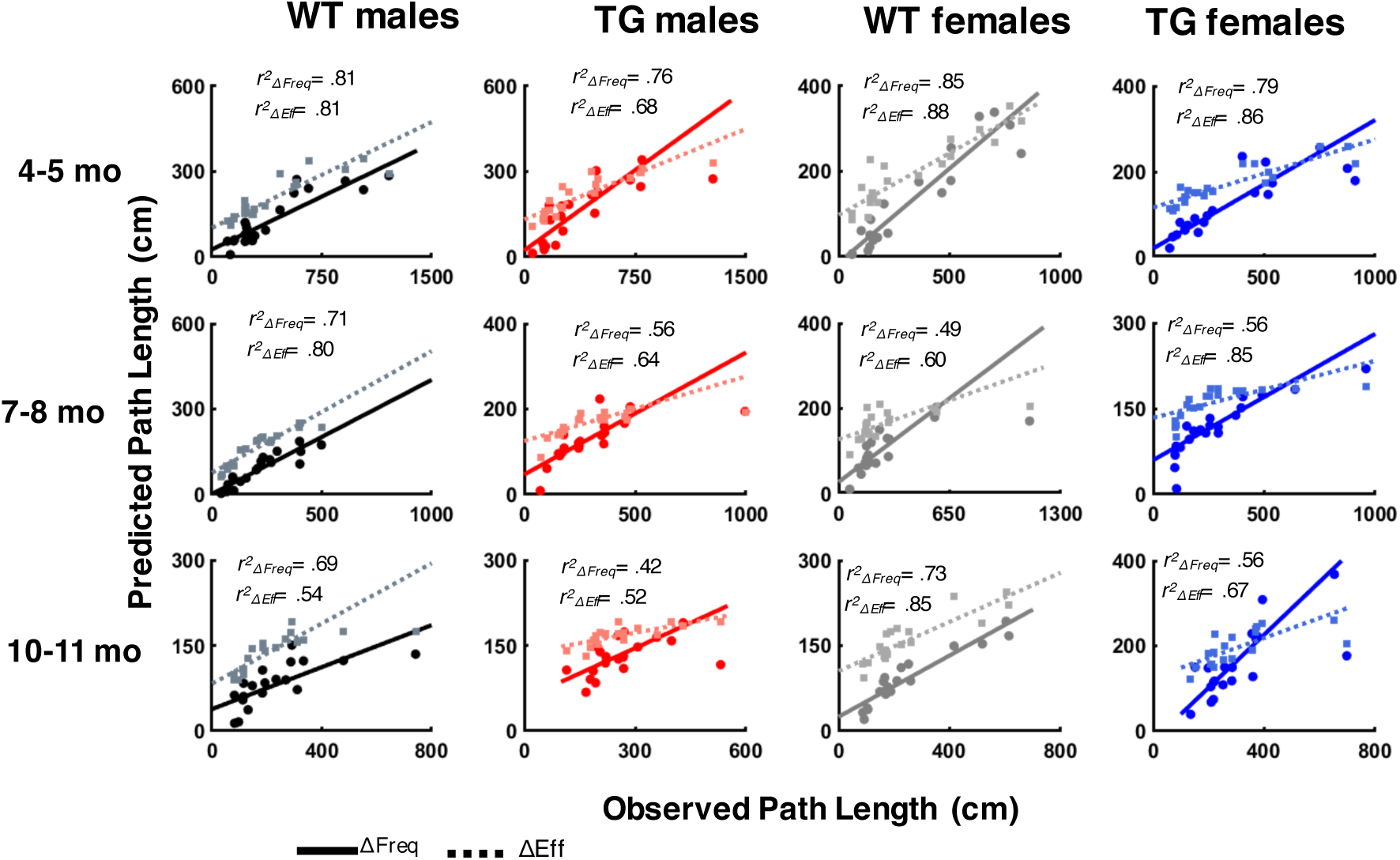
Scatter plots of derived predicted path length (cm) and observed path length (cm). Changing between movements (solid line) and their efficiency (dashed line) was significantly predictive of performance for all animals at each age of testing.

**Figure 9.**
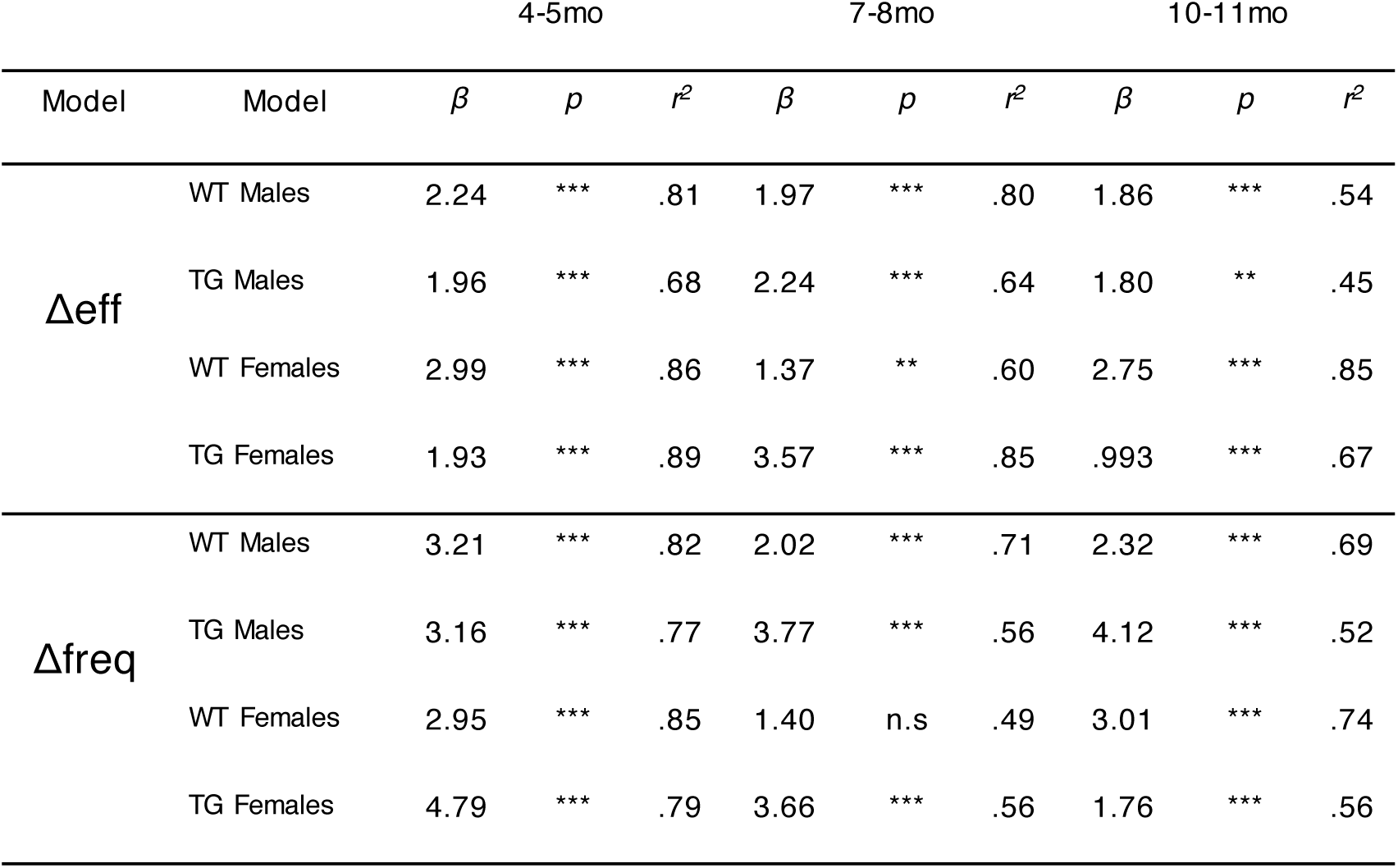
Linear regressions indicate dynamic patterns of swim movements across testing age. Δeff= regression model of predicted score computed by allowing efficiency to modulate over trials (i.e. frequency-fixed). Δfreq = regression model of predicted scores computed allowing frequency to modulate over trials (i.e. efficiency-fixed). *\*\*p*<.01, *\*\*\*p*<.001.

#### Swim speed and Non-Spatial Errors

To determine whether the impairments described above might be influenced by deficits in acquiring the task procedures and sensorimotor behavior, we acquired measurements of swim speed during each training trial (Fig. S1). On average, swim speeds increased as a function of test day within each time-point (*Fs*≥ 4.20, *ps*≤.02). In addition, female rats exhibited faster swim speeds compared male rats—an observation that was consistent at each time-point (*Fs*≥ 15.67, *ps*≤.001). Nevertheless, there were no significant transgenic differences as Tg and WT animals demonstrated similar swim speeds at 10-11 months of age.

We also analyzed the number of non-spatial errors per rat at each age point. The number of errors were summed across days to produce a non-spatial error score (Harker and Whishaw, 2002). Overall, animals in both groups displayed near zero non-spatial error scores at each age of testing (Fig. S2). By 10-11 months of age, only half of the animals in each group expressed 1 or 2 errors over the 5 days of hidden platform testing, indicating an absence of procedural learning deficits in the Tg group (*X^2^(2)=* 7.54, *p*=.37). Thus, given the absence of clear group differences in non-spatial behaviors, it is unlikely that procedural errors contributed to the deficits described in the sections above.

### No-Platform Probe Test

At each testing age, a no-platform probe test was conducted 24 hours after the final day 5 training trial. Figure 10A shows heat maps representing the dwell time in each location of the pool collapsed across animals in each of the Tg and WT groups. The heat maps suggest that rats from each group organized their movements around the trained platform location and spent a disproportionate amount of time near this region. To determine whether the preference for the platform region is expressed for the full duration of the probe session, we divided the analysis into four 15 second bins (Fig. 10B). At each test age, measures of average proximity increased as a function of time bin, suggesting that animals made fewer swims near the platform location by the end of each probe session (*Fs*≥ 3.30, *ps*≤.025). However, measures of target preference score indicated significant differences at 7-8 months of age only, whereby all animals had significantly lower preference for the platform quadrant in the last 15s versus all other time bins (*F*(3,72)=3.132,*p*=.031). Neither tests of proximity nor preference score revealed significant Tg and sex differences or interaction effects at each test age, indicating that groups displayed an equivalent search preference for the platform location by the end of training.

**Figure 10.**
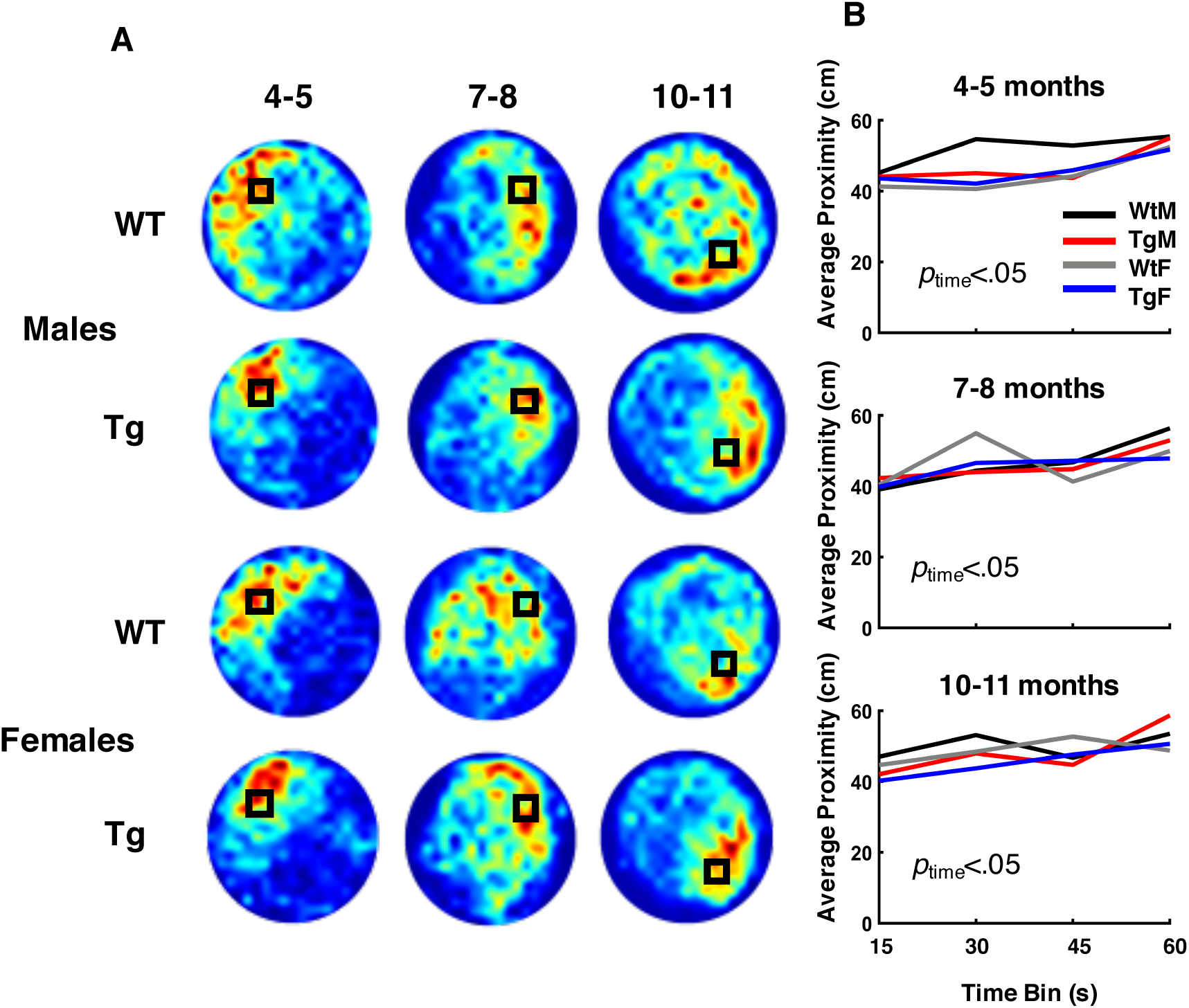
All groups exhibited similar preference for the platform location during the probe trial. (A) Heat maps represent weighted occupancy across entire 60s trial. Hot colors indicate longer dwell times. The platform area is denoted with a black square. (B) Average proximity away from the platform in centimeters is plotted across 15s time bins. All groups demonstrated similar average proximity within each time bin, though proximity to the platform increased as function of time (p<.05, Mixed ANOVA).

### Morris Water Task - Cued Platform

In the cued platform task, Tg and WT rats showed similar performance at each testing age (Fig. S3). Mixed ANOVAs conducted on escape latencies at each time-point indicated all animals had decreased escape latencies in the second trial block versus the first trial block at 4-5 months and 7-5 months (*Fs*≥ 4.46, *ps*≤.0.045), but only trending differences were observed at 10-11 months (*F*(1,23)=3.81, *p*=.063). Importantly, there were no group differences detected between Tg and WT groups at the three test ages. Further, there were no main effects of sex at each testing age. At 4-5 months of age, there was a significant sex by block effect, whereby female animals were slightly slower to reach the platform relative to male animals in the first trial block. Furthermore, Tg females had slightly elevated latencies to reach the platform relative to Tg males, though this genotype by sex effect only trended towards significance (*F*(1,24)=2.97,*p*=.098).

## Discussion

The primary conclusion of the present study is that clear spatial navigation impairments by TgF344-AD rats were identified at 10-11 months of age. Specifically, TgF344-AD rats demonstrated increased escape latencies and path lengths, and they searched a wider area of the pool and were less precise in their search for the platform location (Figs 2 and 3). In addition, by 10-11 months of age, the directionality of their trajectories to the platform (Fig. 5 and 6) and switching from less direct to more direct trajectory types was attenuated in Tg rats (Fig. 8 and 9). While navigation impairments were detected during training in the hidden platform task, a 60 second no-platform probe test conducted on the 6^th^ day indicated that both Tg and WT groups displayed a similar preference for the platform quadrant (Fig. 10). This pattern of impairments at 10-11 months supports the conclusion that although Tg animals are impaired at executing an optimal trajectory and search pattern near the hidden platform region, Tg rats can express a preference for that location by the end of the experiment.

The deficits reported in the present study were observed in the absence of group differences in sensorimotor or procedural learning. Several analyses support this conclusion: first, measures of swim speed failed to indicate significant differences between Tg and WT groups at any age of testing. This observation is also consistent with recent work (Rorabaugh et al., 2017). Secondly, measures of non-spatial errors at each age of testing failed to indicate Tg and WT differences. Lastly, Tg and WT animals performed similarly on the cued platform task at each test age, indicating that Tg animals could learn to navigate by approaching a cue directly marking the goal. These findings strongly suggest that Tg and WT animals were equivalently capable of acquiring the non-spatial demands of the Morris water task.

The results of the present study offer some clarification regarding the time-course of spatial navigation impairment observed in this model. Only a small number of studies have investigated navigation in early stages of pathogenesis in the TgF344-AD model, with one study demonstrating clear deficits in the Morris water task at 6 months of age (Rorabaugh et al., 2017), and others showing largely intact navigation between 4 and 6 months (Cohen et al., 2013; Pentkowski et al., 2017). While the clearest navigation deficits were detected much later at 10-11 months of age in the present study, our results are also consistent with previous work in showing that Tg rats displayed a significant decrease in the directedness of their swim trajectories by 7 months of age. Notably, Tg and WT animals were equivalent in standard measures of water task performance at 7-8 months of age (i.e., escape latency and path length). One possible explanation for the latter finding is that repeated spatial training and procedural learning may have allowed Tg animals to utilize strategies that result in similar performance on standard measures. Indeed, a convolution analysis indicated that Tg rats get better at less direct movements to find the platform than switching to more direct movements. Finally, it is important to note that previous studies report that Tg rats tend to have greater spatial difficulties in manipulations involving “reversal” tests in which the goal is moved to a novel location. Because Tg rats display intact navigation in standard tests at this age range (Cohen et al., 2013; Pentkowski et al., 2017; Rorabaugh et al., 2017), it is possible that reversal impairments may reflect the increasing demands on behavioral flexibility rather than navigation *per se*. This possibility should be explored in future work.

Overall, the spatial impairment in TgF344-AD rats found here also closely correspond to those observed in individuals with preclinical or prodromal AD (Guariglia and Nitrini, 2009; Pai and Jacobs, 2004). Although platform memory remained intact across all time points, Tg subjects exhibited less accurate trajectories and platform search patterns indicative of allocentric navigation impairments. In contrast, the intact cued platform navigation is suggestive of intact egocentric navigation. This pattern of impaired allocentric, and intact egocentric navigation, is consistent with impairments observed in human subjects with early AD (Serino et al., 2014). Thus, it is likely the preclinical phase of TgF344-AD rats lies prior to 10 months, and greater spatial navigation and memory impairment would be observed later. Prior characterization of pathological markers of AD in TgF344-AD rats supports this notion. Specifically, TgF344-AD rats develop neuropathic changes and entorhinal-to-hippocampus synaptic changes as early as 6 months of age (Cohen et al., 2013; Smith and McMahon, 2018) and neurovascular dysfunction and CA3-to-CA1 synaptic changes at 9 months (Joo et al., 2017; Smith and McMahon, 2018), while significant amyloid plaques, inflammation, tauopathy and cell loss occurs at 16 months. Thus, the navigation differences observed at 10-11 months in TgF344-AD males could be considered a putative MCI phase, though further characterization of pathology and behavior is needed.

Although there were no clear differences between male and female Tg rats on measures of path length, platform proximity, or search area, we did observe a trend for greater escape latency at 10-11 months of age. Additionally, our detailed path analyses indicated that female Tg rats performed significantly fewer direct trajectories toward the hidden platform at 10-11 months of age. Interestingly, performance of direct paths by female Tg rats was similar at 7-8 and 10-11 months of age, suggesting a potential 7-month onset of subtle changes in swim path trajectory. Male Tg rats demonstrated a slightly decreased, yet apparent impairment, in direct navigation at 10-11 months. Thus, the attenuated directional deficits found in male Tg rats relative to female Tg rats may reflect sex-specific progression profiles like that found in various models of AD-like pathology (Clinton et al., 2007; Granger et al., 2016; King et al., 1999; Melnikova et al., 2016), and is consistent with human studies in which the prevalence and rate of conversion to AD is higher in females (Gao et al., 1998; Mielke et al., 2014; Pike, 2017) Finally, our observations suggest that detailed path analyses might have greater sensitivity at detecting group differences than general performance measures.

The hippocampus has been a strong focus in AD research, despite various other limbic circuit structural involvement in AD (Aggleton et al., 2016). Cohen and colleagues (2013) identified inflammation and initial tau changes at 6 months of age in the hippocampus, and recent work has shown that this early expression of pathology also impacts the medial entorhinal cortex and entorhinal-hippocampal synaptic plasticity (Rorabaugh et al., 2017; Smith and McMahon, 2018). This coincides with the reported early incidence of pathology in humans associated with the entorhinal cortex (Braak and Braak, 1995), but fails to address whether TgF344-AD pathology emerges in other limbic regions such as the anterior thalamic nuclei or retrosplenial cortex. This is particularly important given cell types coding for head direction are found in both regions (Clark and Taube, 2012; Taube, 2007), and damage to both regions can produce deficits in the directional accuracy of navigation (Clark and Harvey, 2016; Harvey et al., 2017; Vann et al., 2009). Whether TgF344-AD rats exhibit AD pathology and disrupted spatial signaling in limbic-thalamic and limbic-cortical regions at early stages of development is unknown and warrants investigation.

### Conclusions

The present study found that TgF344-AD rats express clear navigation impairments at 10-11 months of age. A detailed path analysis indicated that subtle deficits in the directedness of trajectories to the hidden platform can be detected at earlier ages and can be sensitive to sex differences. The latter observations may underlie the subtle differences between Tg and WT rats that were found using classic measures of water maze (escape latency, path length, search proximity and search area) at these ages, and are likely not due to factors associated with non-spatial task demands. Furthermore, spatial memory was intact for all animals across all ages indicating that developing deficits observed at 10-11 months in Tg animals may be indicative of an MCI stage of disease progression. Future work should identify whether the observed navigational impairments in this model map onto brain regions involved in directional computation, such as the anterior thalamus or retrosplenial cortex. Overall, the TgF344-AD rat model provides substantial promise for elucidating neurobiological mechanisms of spatial disorientation in AD.

## Acknowledgments

This research was supported by grants from the Alzheimer’s Association (AARG-17-531572), NIGMS (P30GM103400), and an NM-INBRE (P20GM10345). The authors thank Elizabeth Sneddon, John Madden, and Sara Benthem for assistance with behavioral analysis and Dr. Derek Hamilton for technical support.

---

**Figure S1.**
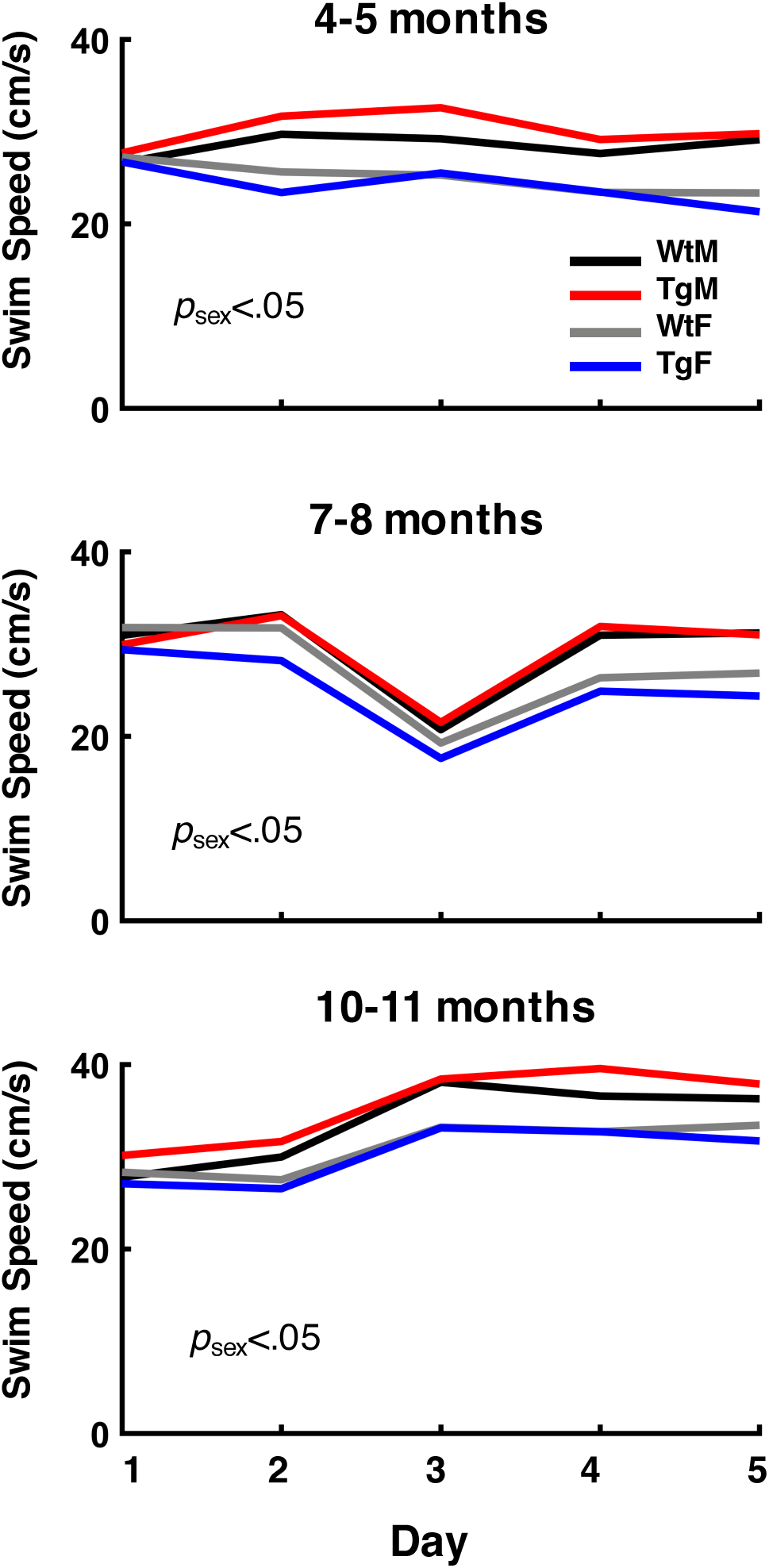
Average swim speed (cm/s) is plotted for each age of testing. Groups distinguished by color: black=Wild Type males (WtM), red=Transgenic males (TgM), gray=Wild Type females (WtF), blue=Transgenic females (TgF). Note that males consistently swam faster than females across all age points (ps<.05, ANOVAs).

**Figure S2.**
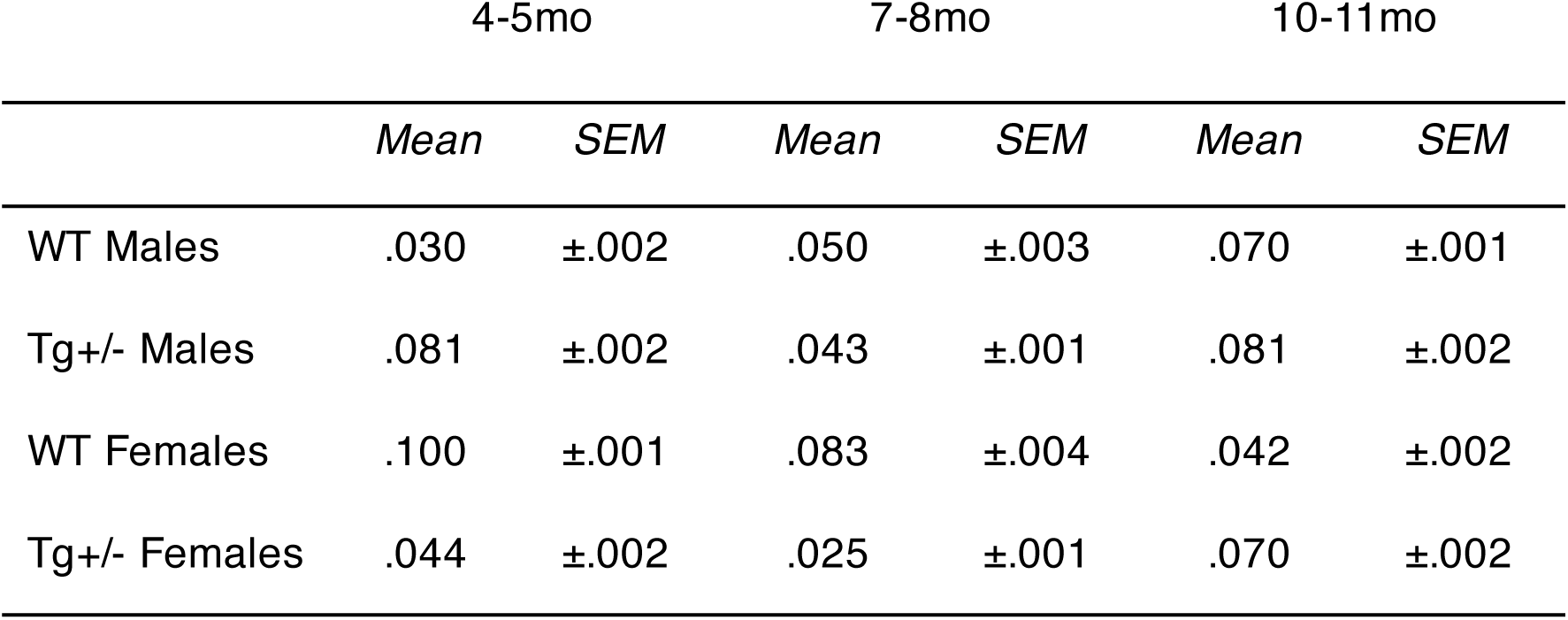
Table showing the means and SEM on non-spatial errors at each age of testing

**Figure S3.**
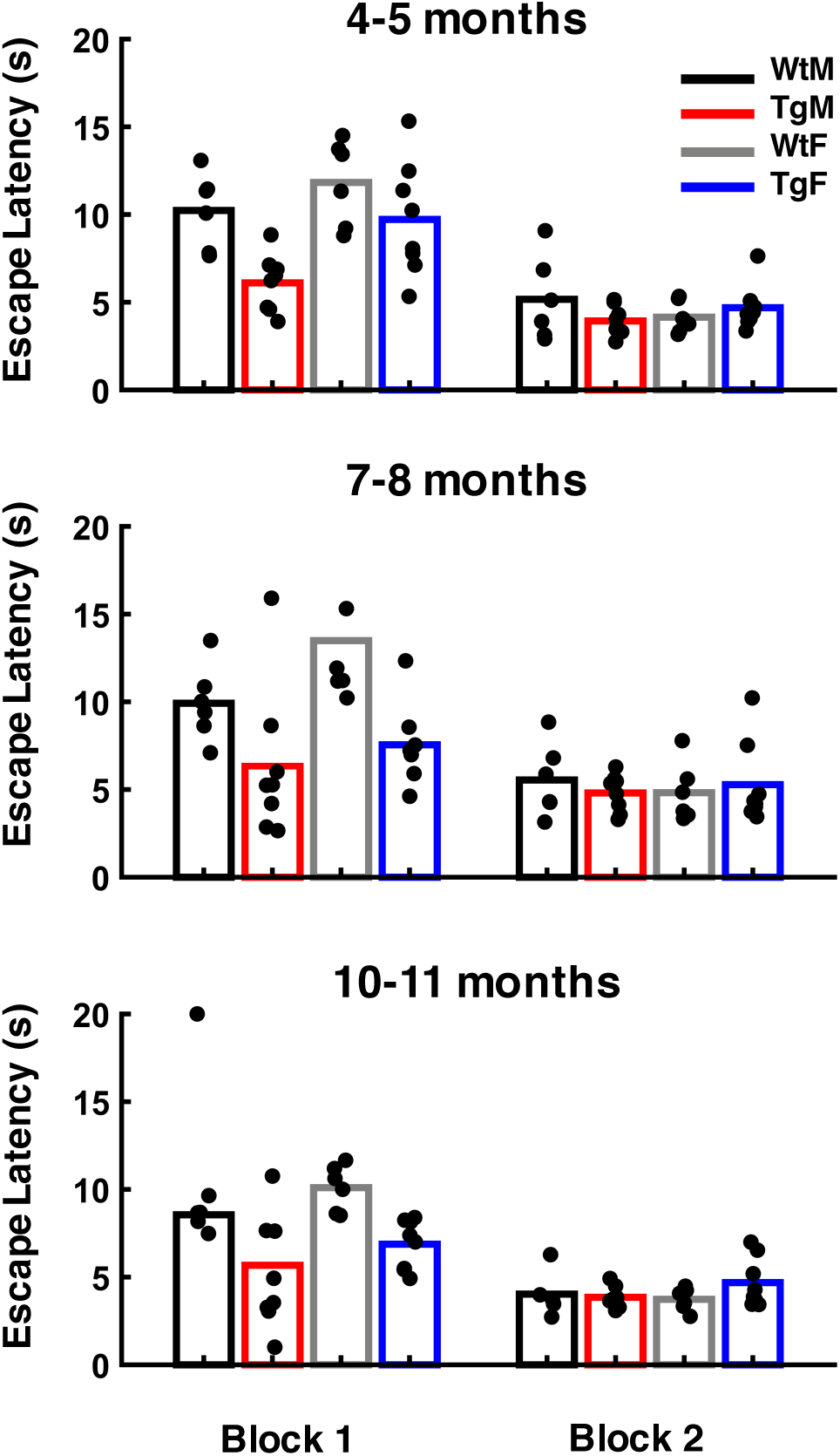
Mean escape latency (s) across trial blocks (set of 4 trials per block) during cued task was not different between groups at each age of testing. Mean latencies are plotted for each animal (black circles) to show distribution. Groups distinguished by color: black=Wild Type males (WtM), red=Transgenic males (TgM), gray=Wild Type females (WtF), blue=Transgenic females (TgF).

